# Subcortical brain volume, regional cortical thickness and cortical surface area across attention-deficit/hyperactivity disorder (ADHD), autism spectrum disorder (ASD), and obsessive-compulsive disorder (OCD)

**DOI:** 10.1101/673012

**Authors:** Premika S.W. Boedhoe, Daan van Rooij, Martine Hoogman, Jos W.R. Twisk, Lianne Schmaal, Yoshinari Abe, Pino Alonso, Stephanie H. Ameis, Anatoly Anikin, Alan Anticevic, Philip Aherson, Celso Arango, Paul D. Arnold, Francesca Assogna, Guillaume Auzias, Tobias Banaschewski, Alexander Baranov, Marcelo C. Batistuzzo, Sarah Baumeister, Ramona Baur-Streubel, Marlene Behrmann, Mark A. Bellgrove, Francesco Benedetti, Jan C. Beucke, Joseph Biederman, Irene Bollettini, Anushree Bose, Janita Bralten, Ivanei E. Bramati, Daniel Brandeis, Silvia Brem, Brian P. Brennan, Geraldo F. Busatto, Sara Calderoni, Anna Calvo, Rosa Calvo, Francisco X. Castellanos, Mara Cercignani, Tiffany M. Chaim-Avancini, Kaylita C. Chantiluke, Yuqi Cheng, Kang Ik K. Cho, Anastasia Christakou, David Coghill, Annette Conzelmann, Ana I. Cubillo, Anders M. Dale, Sara Dallaspezia, Eileen Daly, Damiaan Denys, Christine Deruelle, Adriana Di Martino, Ilan Dinstein, Alysa E. Doyle, Sarah Durston, Eric A. Earl, Christine Ecker, Stefan Ehrlich, Benjamin A. Ely, Jeffery N. Epstein, Thomas Ethofer, Damien A. Fair, Andreas J. Fallgatter, Stephen V. Faraone, Jennifer Fedor, Xin Feng, Jamie D. Feusner, Jackie Fitzgerald, Kate D. Fitzgerald, Jean-Paul Fouche, Christine M. Freitag, Egill A. Fridgeirsson, Thomas Frodl, Matt C. Gabel, Louise Gallagher, Tinatin Gogberashvili, Ilaria Gori, Patricia Gruner, Deniz A. Gürsel, Shlomi Haar, Jan Haavik, Geoffrey B. Hall, Neil A. Harrison, Catharina A. Hartman, Dirk J. Heslenfeld, Yoshiyuki Hirano, Pieter J. Hoekstra, Marcelo Q. Hoexter, Sarah Hohmann, Marie F. Høvik, Hao Hu, Chaim Huyser, Neda Jahanshad, Maria Jalbrzikowski, Anthony James, Joost Janssen, Fern Jaspers-Fayer, Terry L. Jernigan, Dmitry Kapilushniy, Bernd Kardatzki, Georgii Karkashadze, Norbert Kathmann, Christian Kaufmann, Clare Kelly, Sabin Khadka, Joseph A. King, Kathrin Koch, Gregor Kohls, Kerstin Kohls, Masaru Kuno, Jonna Kuntsi, Gerd Kvale, Jun Soo Kwon, Luisa Lázaro, Sara Lera-Miguel, Klaus-Peter Lesch, Liesbeth Hoekstra, Yanni Liu, Christine Lochner, Mario R. Louza, Beatriz Luna, Astri J. Lundervold, Charles B. Malpas, Paulo Marques, Rachel Marsh, Ignacio Martínez-Zalacaín, David Mataix-Cols, Paulo Mattos, Hazel McCarthy, Jane McGrath, Mitul A. Mehta, José M. Menchón, Maarten Mennes, Mauricio Moller Martinho, Pedro S. Moreira, Astrid Morer, Pedro Morgado, Filippo Muratori, Clodagh M. Murphy, Declan G.M. Murphy, Akiko Nakagawa, Takashi Nakamae, Tomohiro Nakao, Leyla Namazova-Baranova, Janardhanan. C. Narayanaswamy, Rosa Nicolau, Joel T. Nigg, Stephanie E. Novotny, Erika L. Nurmi, Eileen Oberwelland Weiss, Ruth L. O’Gorman Tuura, Kirsten O’Hearn, Joseph O’Neill, Jaap Oosterlaan, Bob Oranje, Yannis Paloyelis, Mara Parellada, Paul Pauli, Chris Perriello, John Piacentini, Fabrizio Piras, Federica Piras, Kerstin J. Plessen, Olga Puig, J. Antoni Ramos-Quiroga, Y.C. Janardhan Reddy, Andreas Reif, Liesbeth Reneman, Alessandra Retico, Pedro G.P. Rosa, Katya Rubia, Oana Georgiana Rus, Yuki Sakai, Anouk Schrantee, Lena Schwarz, Lizanne J.S. Schweren, Jochen Seitz, Philip Shaw, Devon Shook, Tim J. Silk, H. Blair Simpson, Norbert Skokauskas, Juan Carlos Soliva Vila, Anastasia Solovieva, Noam Soreni, Carles Soriano-Mas, Gianfranco Spalletta, Emily R. Stern, Michael C. Stevens, S. Evelyn Stewart, Gustavo Sudre, Philip R. Szeszko, Leanne Tamm, Margot J. Taylor, David F. Tolin, Michela Tosetti, Fernanda Tovar-Moll, Aki Tsuchiyagaito, Theo G.M. van Erp, Guido A. van Wingen, Alasdair Vance, Ganesan Venkatasubramanian, Oscar Vilarroya, Yolanda Vives-Gilabert, Georg G. von Polier, Susanne Walitza, Gregory L. Wallace, Zhen Wang, Thomas Wolfers, Yuliya N. Yoncheva, Je-Yeon Yun, Marcus V. Zanetti, Fengfeng Zhou, Georg C. Ziegler, Kathrin C. Zierhut, Marcel P. Zwiers, the ENIGMA-ADHD working group, the ENIGMA-ASD working group, the ENIGMA-OCD working group, Paul M. Thompson, Dan J. Stein, Jan Buitelaar, Barbara Franke, Odile A. van den Heuvel

**Author notes:** Please cc. as corresponding author: prof.dr. O.A. van den Heuvel. equal contribution (Daan van Rooij, Martine Hoogman). see excel sheet for full list of authors (the ENIGMA-ADHD working group, the ENIGMA-ASD working group, the ENIGMA-OCD working group). Location of work and address for reprints: Premika S.W. Boedhoe, M.Sc., Department of Psychiatry, Amsterdam UMC Location VUmc, PO Box 7057, 1007 MB, Amsterdam, The Netherlands.

## Abstract

**Objective:** Attention-deficit/hyperactivity disorder (ADHD), autism spectrum disorder (ASD), and obsessive-compulsive disorder (OCD) are common neurodevelopmental disorders that frequently co-occur. We aimed to directly compare all three disorders. The ENIGMA consortium is ideally positioned to investigate structural brain alterations across these disorders.

**Methods:** Structural T1-weighted whole-brain MRI of controls (*n*=5,827) and patients with ADHD (n=2,271), ASD (n=1,777), and OCD (n=2,323) from 151 cohorts worldwide were analyzed using standardized processing protocols. We examined subcortical volume, cortical thickness and surface area differences within a mega-analytical framework, pooling measures extracted from each cohort. Analyses were performed separately for children, adolescents, and adults using linear mixed-effects models adjusting for age, sex and site (and ICV for subcortical and surface area measures).

**Results:** We found no shared alterations among all three disorders, while shared alterations between any two disorders did not survive multiple comparisons correction. Children with ADHD compared to those with OCD had smaller hippocampal volumes, possibly influenced by IQ. Children and adolescents with ADHD also had smaller ICV than controls and those with OCD or ASD. Adults with ASD showed thicker frontal cortices compared to adult controls and other clinical groups. No OCD-specific alterations across different age-groups and surface area alterations among all disorders in childhood and adulthood were observed.

**Conclusion:** Our findings suggest robust but subtle alterations across different age-groups among ADHD, ASD, and OCD. ADHD-specific ICV and hippocampal alterations in children and adolescents, and ASD-specific cortical thickness alterations in the frontal cortex in adults support previous work emphasizing neurodevelopmental alterations in these disorders.

## Introduction

Attention-deficit/hyperactivity disorder (ADHD), autism spectrum disorder (ASD), and obsessive-compulsive disorder (OCD) are common neurodevelopmental disorders with a lifetime prevalence of 2.5-5%, ∼1%, and 2.3%, respectively (1-3). Symptoms mostly develop early in life (ADHD, ASD) or later in childhood (OCD) and often persist into adulthood. Characteristic symptoms include inattentiveness, impulsivity and hyperactivity for ADHD; impairments in social communication and restricted and stereotyped behaviors for ASD; and repetitive thoughts (obsessions) and behaviors or mental acts (compulsions) that cause distress or anxiety for OCD. Although each disorder is distinguished by its own core symptoms, the disorders frequently co-occur and share overlap in phenomenology and pathophysiology (4,5).

There are parallels between the uncontrollable impulsive behaviors of ADHD and the excessive and compulsive rituals of OCD and ASD. Impaired response inhibition and cognitive control processes may underlie the cross-disorder traits within the impulsive-compulsive spectrum (6), implicating cortico-striato-thalamo-cortical and fronto-parietal networks (7). It remains unclear which morphological brain abnormalities within these networks are shared (non-specific) versus distinct (specific to one disorder).

Imaging studies, including meta-analyses, have generally compared one of the three disorders to healthy controls (8-12). Large-scale studies generally yielded small to moderate effect sizes, indicating that disorder-associated alterations are subtle (13-17). Few structural imaging studies have directly compared these three disorders (18,19), mostly in small numbers and with inconsistent results (20). A meta-analysis including 931 patients with ADHD and 928 with OCD reported shared smaller ventromedial prefrontal cortex gray matter volume, ADHD-specific smaller gray matter volume in basal ganglia and insula, and OCD-specific smaller volume of rostral and dorsal anterior cingulate and medial prefrontal cortex (21). Another meta-analysis comparing structural brain alterations in 911 patients with ASD and 928 with OCD reported shared alterations in the dorsal medial prefrontal cortex and OCD-specific alterations in the basal ganglia (22). However, despite their clinical overlap, no structural gray matter study so far compared all three disorders.

The ENIGMA consortium (23) includes the largest samples for ADHD, ASD, and OCD worldwide (13-17). The consortium also improves on earlier meta-analyses by using harmonized protocols for brain segmentation and quality control procedures across ENIGMA working groups, and by pooling extracted individual participant data. The ENIGMA consortium is therefore ideally positioned to investigate overlap and specificity of structural brain alterations across disorders.

Here, we present the largest comparative study investigating subcortical and cortical alterations across ADHD, ASD, and OCD. We extracted subcortical volumes, cortical thickness, and cortical surface area estimates of 12,198 individuals from 151 cohorts worldwide, using harmonized data processing protocols. Based on previous meta- and mega-analyses, we expected to find ADHD-specific alterations in frontal and temporal surface areas and basal ganglia volumes in children (14,15), ASD-specific alterations in frontal and temporal cortical thickness (13), and OCD-specific alterations in the thalamus of pediatric patients and the pallidum of adult patients (16). We expected that alterations in the striatum and dorsomedial prefrontal cortex would be observed across disorders (21,22).

## Methods

### Samples

The ENIGMA-ADHD working group includes 48 cohorts from 34 research institutes, with neuroimaging and clinical data from patients with ADHD and healthy controls. The ENIGMA-ASD working group includes 56 cohorts from 38 research institutes, with neuroimaging and clinical data from patients with ASD and healthy controls. The ENIGMA-OCD working group includes 47 cohorts from 34 research institutes, with neuroimaging and clinical data from patients with OCD and healthy controls (Figure 1).

**Figure 1:**
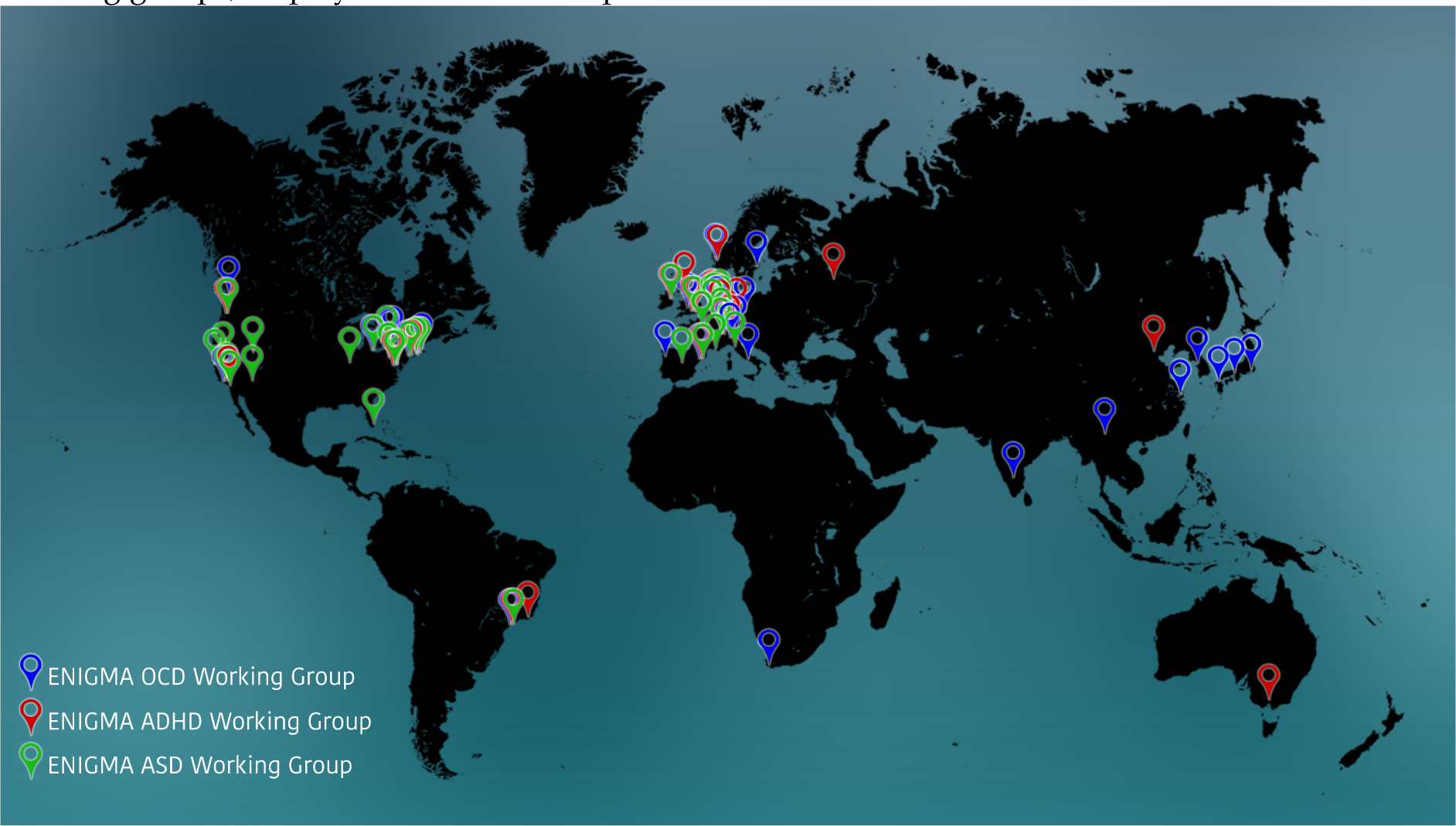
Overview of research institutes participating in the ENIGMA ADHD, ASD and OCD working groups, displayed on a world map

All working groups included data from subjects across the lifespan. As prior results suggested differential effects between pediatric (<12 years), adolescent (12-18 years), and adult (≥18 years) patients, we performed separate mega-analyses for these three age-groups. In total, we analyzed data from 2,271 patients with ADHD, 1,777 with ASD, 2,323 with OCD, and 5,827 healthy controls. All local institutional review boards permitted the use of measures extracted from the coded data for mega-analyses.

### Image Acquisition and Processing

Structural T_1_-weighted whole-brain MRI was acquired and processed locally. Image acquisition parameters for each cohort are listed in Supplementary Tables S1-S3. All cortical parcellations were performed with the fully automated segmentation program FreeSurfer, version 5.3, following standardized ENIGMA protocols to harmonize analyses and quality control procedures across multiple sites (http://enigma.ini.usc.edu/protocols/imaging-protocols/). Segmentations of seven bilateral subcortical and 34 bilateral cortical regions of interest according to the Desikan-Killiany atlas were visually inspected and statistically evaluated for outliers. Details on image exclusion criteria and quality control are presented in Supplementary Information SI1. All cohorts of each working group underwent identical processing and quality control procedures.

### Statistical Analysis

We pooled extracted subcortical volumes, cortical thickness and cortical surface area measures from individual subjects across all cohorts from the different working groups into one database to perform a mega-analysis. We examined differences among patients groups and controls using linear mixed-effects models in STATA; mixed models are used to take into account the differences between sites. The means of the left and right hemisphere of 34 cortical regions (separately for cortical thickness and cortical surface area), whole-hemisphere measures (average thickness and total surface area), and seven subcortical regions were used in the mega-analyses. To obtain comparable standardized regression coefficients (effect sizes) for all comparisons the z-scores for each of the cortical and subcortical regions-of-interest served as the outcome measures, and the diagnoses (ADHD, ASD, OCD, and HC) were included as independent variables of interest, using three dummy variables. Disorder specific alterations were assessed by alternating the different diagnoses as reference category. Shared alterations were assessed using the HC as a reference category. A random intercept for cohort was entered to account for clustering within cohorts; if necessary (i.e. when there was a significant improvement of the model fit), a random slope for diagnosis*cohort was included to account for different effect sizes between cohorts within the different working groups (24). Age and sex were included as covariates (25,26); for the surface area and subcortical volume analyses, ICV was also added as a covariate, since these measures scale with head size (27).

To detect potentially different effects of disorder with age, we performed all analyses separately for pediatric, adolescent, and adult patients. Because only a limited number of cohorts had data on IQ and medication use, sensitivity analyses were performed to investigate how IQ and psychotropic medication use might have influenced the disorder differences. For medication use (yes/no at time of scanning), stratified analyses according to medication status were performed. With respect to IQ, we included the variable as an additional covariate in the analyses. The Benjamini-Hochberg false discovery rate (FDR) was used to control for multiple comparisons, for 35 cortical measures (separately for cortical thickness and surface area) and 7 subcortical measures, and standardized effect sizes were obtained. Results were considered significant if the FDR-corrected p-value (q) was ≤0.05.

## Results

The demographic and clinical characteristics of participants are summarized per age category in Table 1a-c (entire sample Supplementary Table S4). Results not surviving multiple comparison correction, but with p-values <0.05 were considered trends and are described for the main analyses in Supplemental Information SI2.

**Table 1a:** demographics, clinical characteristics, age, sex, and numbers breakdown separately for pediatric patient roups and control subjects

| children |  | OCD (N total patients=140) |  |  | ADHD (N total patients=709) |  |  | ASD (N total patients=723) |  |  | controls (N total controls=1590) |  |  |
| --- | --- | --- | --- | --- | --- | --- | --- | --- | --- | --- | --- | --- | --- |
|  |  | number of sites included (14) |  |  | number of sites included (26) |  |  | number of sites included (35) |  |  | number of sites included (69) |  |  |
|  |  | data available for N patients | mean | sd | data available for N patients | mean | sd | data available for N patients | mean | sd | data available for N subjects | mean | sd |
| age |  | 140 | 10.28 | 1.22 | 709 | 9.41 | 1.33 | 723 | 8.64 | 2.46 | 1590 | 9.35 | 1.72 |
| IQ |  | 70 | 108.75 | 16.50 | 648 | 106.09 | 15.20 | 526 | 100.02 | 21.49 | 1302 | 111.23 | 15.37 |
|  |  | data available for N patients | N | % | data available for N patients | N | % | data available for N patients | N | % | data available for N subjects | N | % |
| male |  | 140 | 76 | 54.29 | 709 | 530 | 74.75 | 723 | 588 | 81.33 | 1590 | 997 | 62.70 |
| medication |  | 140 | 38 | 27.14 | 438 | 126 | 28.77 | 413 | 97 | 23.49 | NA |  |  |
| comorbid | OCD | - | - | - | 408 | 0 | 0.00 | 73 | 0 | 0.00 |  |  |  |
|  | ADHD | 126 | 14 | 11.11 | - | - | - | 73 | 8 | 10.96 |  |  |  |
|  | ASD | 126 | 3 | 2.38 | 271 | 0 | 0.00 | - | - | - |  |  |  |
|  | TD | 119 | 12 | 10.08 | 240 | 1 | 0.42 | 73 | 0 | 0.00 |  |  |  |
|  | Anx | 129 | 41 | 31.78 | 408 | 21 | 5.15 | 73 | 4 | 5.48 |  |  |  |
|  | Dep | 129 | 7 | 5.43 | 404 | 0 | 0.00 | 73 | 1 | 1.37 |  |  |  |

Abbreviations: sd= standard deviation; TD = Tourette's Disorder; Anx = Anxiety Disorder; Dep= Major Depressive Disorder

**Table 1b:** demographics, clinical characteristics, age, sex, and numbers breakdown separately for adolescent Patient groups and control subjects

| adolescents |  | OCD (N total patients=359) |  |  | ADHD (N total patients=633) |  |  | ASD (N total patients=565) |  |  | controls (N total controls=1368) |  |  |
| --- | --- | --- | --- | --- | --- | --- | --- | --- | --- | --- | --- | --- | --- |
|  |  | number of sites included (16) |  |  | number of sites included (27) |  |  | number of sites included (39) |  |  | number of sites included (79) |  |  |
|  |  | data available for N patients | mean | sd | data available for N patients | mean | sd | data available for N patients | mean | sd | data available for N subjects | mean | sd |
| age |  | 359 | 14.91 | 1.72 | 633 | 14.00 | 1.65 | 565 | 14.40 | 1.66 | 1368 | 14.37 | 1.71 |
| IQ |  | 136 | 106.75 | 14.31 | 608 | 102.41 | 14.63 | 467 | 103.02 | 18.26 | 1085 | 110.07 | 12.79 |
|  |  | data available for N patients | N | % | data available for N patients | N | % | data available for N patients | N | % | data available for N subjects | N | % |
| male |  | 359 | 195 | 54.32 | 633 | 514 | 81.20 | 565 | 492 | 87.08 | 1368 | 937 | 68.49 |
| medication |  | 357 | 172 | 48.18 | 487 | 212 | 43.53 | 227 | 74 | 32.60 | NA |  |  |
| comorbid | OCD | - | - | - | 452 | 0 | 0.00 | 68 | 0 | 0.00 |  |  |  |
|  | ADHD | 314 | 27 | 8.60 | - | - | - | 68 | 13 | 19.12 |  |  |  |
|  | ASD | 299 | 10 | 3.34 | 316 | 7 | 2.22 | - | - | - |  |  |  |
|  | TD | 314 | 17 | 5.41 | 272 | 1 | 0.37 | 68 | 1 | 1.47 |  |  |  |
|  | Anx | 316 | 109 | 34.49 | 452 | 8 | 1.77 | 68 | 3 | 4.41 |  |  |  |
|  | Dep | 317 | 24 | 7.57 | 452 | 5 | 1.11 | 68 | 2 | 2.94 |  |  |  |

Abbreviations: sd= standard deviation; TD = Tourette's Disorder; Anx = Anxiety Disorder; Dep= Major Depressive Disorder

**Table 1c:** demographics, clinical characteristics, age, sex, and numbers breakdown separately for adult patient roups and control subjects

| adults |  | OCD (N total patients=1824) |  |  | ADHD (N total patients=929) |  |  | ASD (N total patients=489) |  |  | controls (N total controls=2869) |  |  |
| --- | --- | --- | --- | --- | --- | --- | --- | --- | --- | --- | --- | --- | --- |
|  |  | number of sites included (33) |  |  | number of sites included (28) |  |  | number of sites included (36) |  |  | number of sites included (91) |  |  |
|  |  | data available for N patients | mean | sd | data available for N patients | mean | sd | data available for N patients | mean | sd | data available for N subjects | mean | sd |
| age |  | 1824 | 31.69 | 9.66 | 929 | 29.82 | 10.19 | 489 | 26.03 | 9.00 | 2869 | 29.74 | 9.88 |
| IQ |  | 408 | 105.99 | 13.22 | 765 | 106.91 | 14.52 | 439 | 108.96 | 16.01 | 1449 | 112.20 | 13.36 |
|  |  | data available for N patients | N | % | data available for N patients | N | % | data available for N patients | N | % | data available for N subjects | N | % |
| male |  | 1824 | 923 | 50.60 | 929 | 622 | 66.95 | 489 | 432 | 88.34 | 2869 | 1634 | 56.95 |
| medication |  | 1803 | 829 | 45.98 | 650 | 119 | 18.31 | 226 | 46 | 20.35 | NA |  |  |
| comorbid | OCD | - | - | - | 787 | 3 | 0.38 | 116 | 3 | 2.59 |  |  |  |
|  | ADHD | 1142 | 51 | 4.47 | - | - | - | 116 | 3 | 2.59 |  |  |  |
|  | ASD | 1079 | 1 | 0.09 | 343 | 0 | 0.00 | - | - | - |  |  |  |
|  | TD | 1182 | 22 | 1.86 | 389 | 4 | 1.03 | 116 | 0 | 0.00 |  |  |  |
|  | Anx | 1491 | 276 | 18.51 | 768 | 13 | 1.69 | 116 | 0 | 0.00 |  |  |  |
|  | Dep | 1513 | 193 | 12.76 | 731 | 7 | 0.96 | 116 | 6 | 5.17 |  |  |  |

Abbreviations: sd= standard deviation; TD = Tourette's Disorder; Anx = Anxiety Disorder; Dep= Major Depressive Disorder

### Shared subcortical and cortical alterations across clinical groups compared to healthy controls

#### Children

with ADHD and those with ASD showed some overlap in *subcortical volume* and *cortical thickness* alterations compared to controls (Supplementary Information SI2), however none of these results survived multiple comparison correction (Supplementary Tables S5-S6). In *<u>adolescents</u>*, we did not observe shared *subcortical and cortical* alterations among any of the disorders (Supplementary Tables S7-S9). *<u>Adult</u>* patients with OCD and those with ASD showed smaller hippocampal volumes compared to adult controls, however this finding did not survive multiple comparison correction in adults with ASD (Supplementary Table S10). Adult patient groups showed no overlap in *cortical* alterations (Supplementary Tables S11-S12). Details on alterations compared to healthy controls per patient group can be found in Supplementary Tables S5-S13.

### Disease-specific subcortical and cortical alterations

#### Children

Figure 2a depicts the pattern of *subcortical volume* alterations in children. Children with ADHD showed significantly smaller ICV compared to those with ASD (effect size=-0.23) or OCD (effect size=-0.28). Children with ADHD (effect size=-0.22) also showed smaller hippocampal volumes compared to children with OCD. No significant *cortical* differences among disorders survived multiple comparison correction (Supplementary Tables S15-S16 & Supplementary Information SI2).

**Figure 2a:**
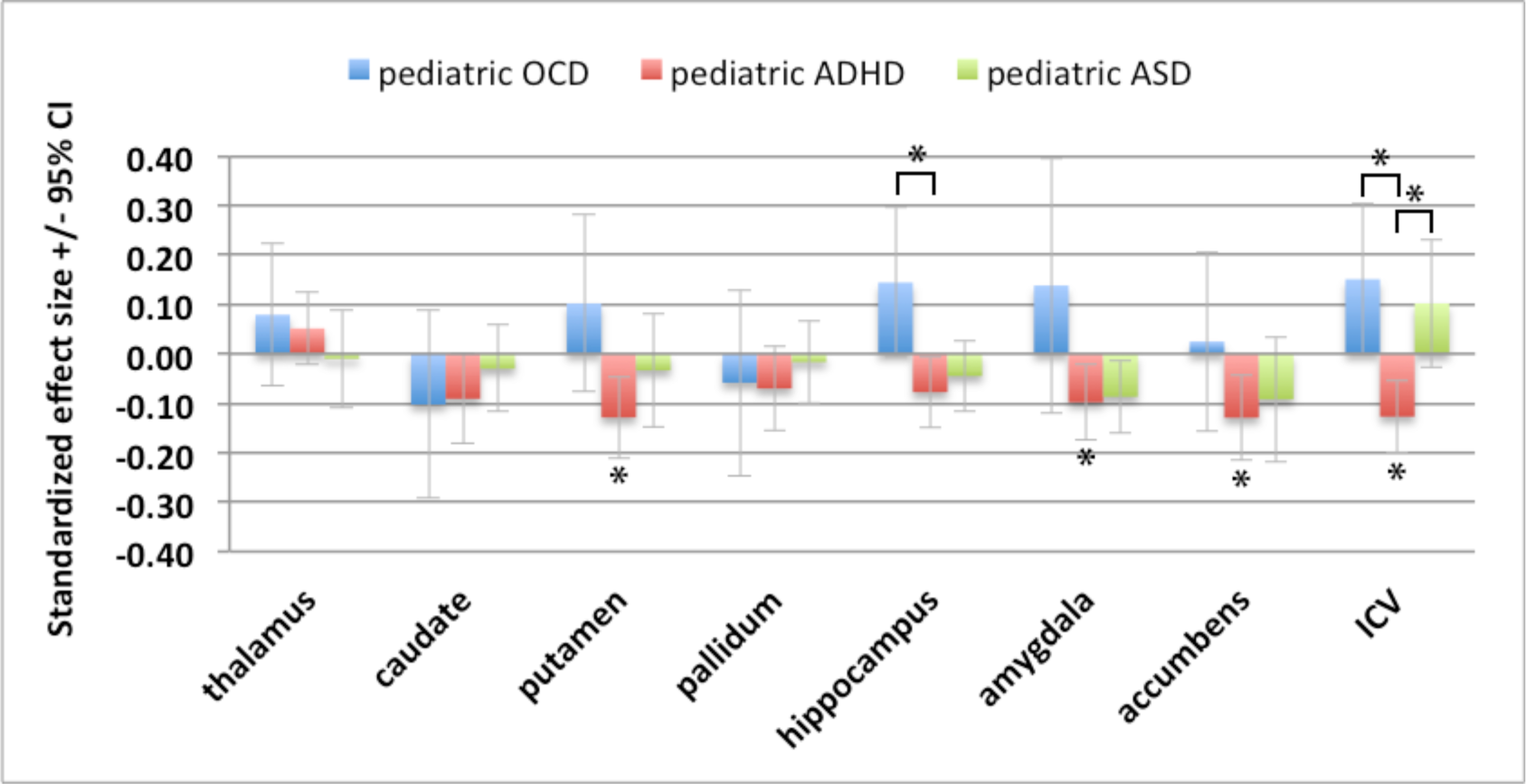
Subcortical volume differences in children with ADHD, ASD, or OCD compared to controls. Significant results (FDR q ≤ 0.05) are indicated by an asterisk (Supplementary Table S5). For Effect size values across disorders see Supplementary Table S14. Abbreviations: Confidence Interval (CI); Intracranial volume (ICV)

#### Adolescents

Adolescents with ADHD had significantly smaller ICV compared to those with ASD (effect size*=*-0.22) or OCD (effect size=-0.19), shown in Figure 2b (Supplementary Table S17). However, the latter did not survive multiple comparison correction. Group differences in *cortical thickness* did not survive multiple comparison correction (Supplementary Table S18 & Supplementary Information SI2). *Surface area* analysis revealed significantly lower surface area of the medial orbitofrontal cortex (effect size=-0.22) in patients with OCD compared to patients with ADHD (Supplementary Table S19).

**Figure 2b:**
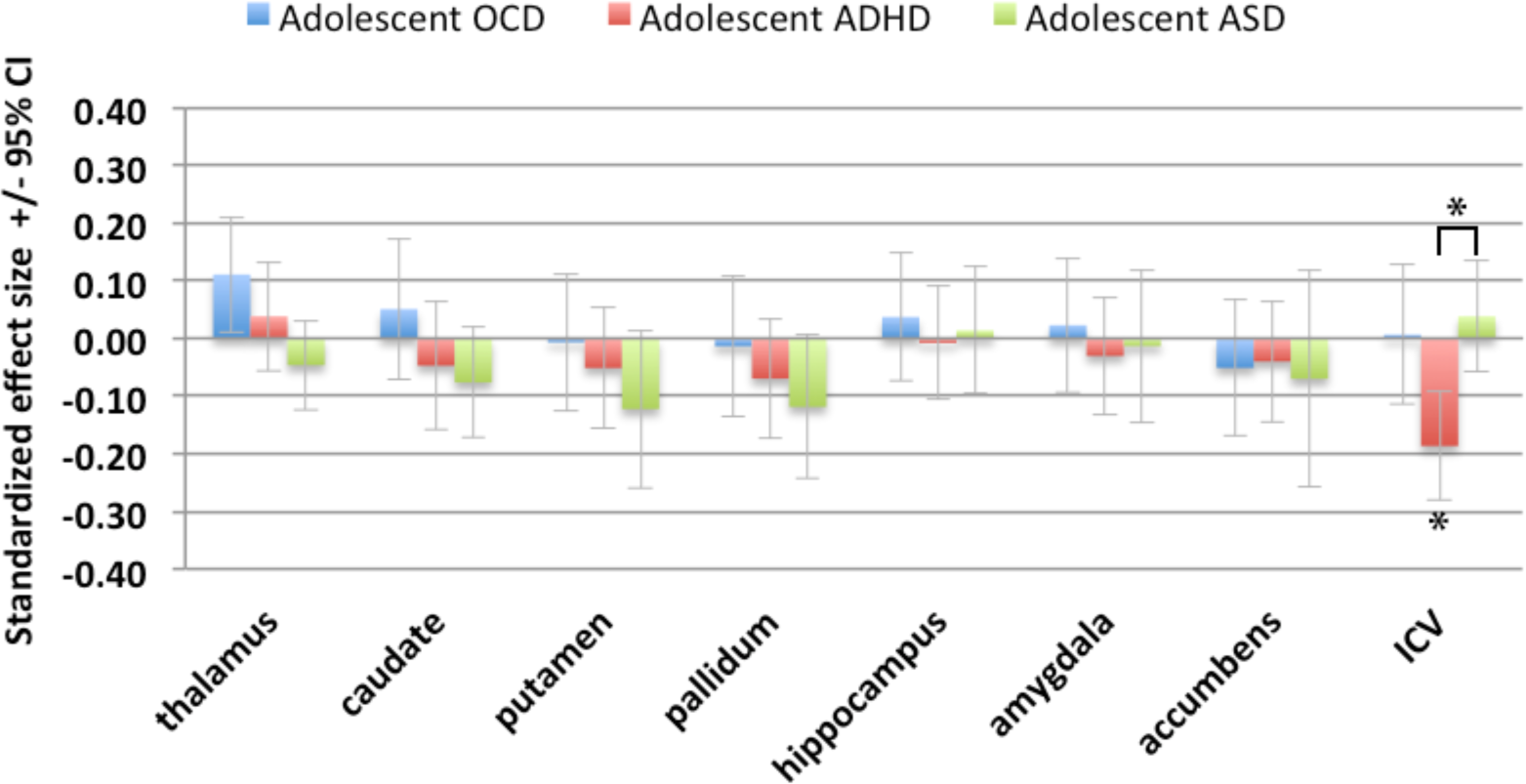
Subcortical volume differences in adolescents with ADHD, ASD, or OCD compared to controls. Significant results (FDR q ≤ 0.05) are indicated by an asterisk (Supplementary Table S7). For Effect size values across disorders see Supplementary Table S17. Abbreviations: Confidence Interval (CI); Intracranial volume (ICV)

#### Adults

None of the *subcortical* volumes differed significantly among adult patient groups (Figure 2c & Supplementary Table S20). *Cortical thickness* analysis revealed significantly thicker cortical gray matter in several frontal regions in adults with ASD compared to adults with OCD or ADHD (Figure 3) with effect sizes varying between 0.17 and 0.30. Adults with OCD did not differ significantly from those with ADHD (Supplementary Table S21). *Surface area* analysis revealed that none of the regions differed significantly among patient groups (Supplementary Table S22).

**Figure 2c:**
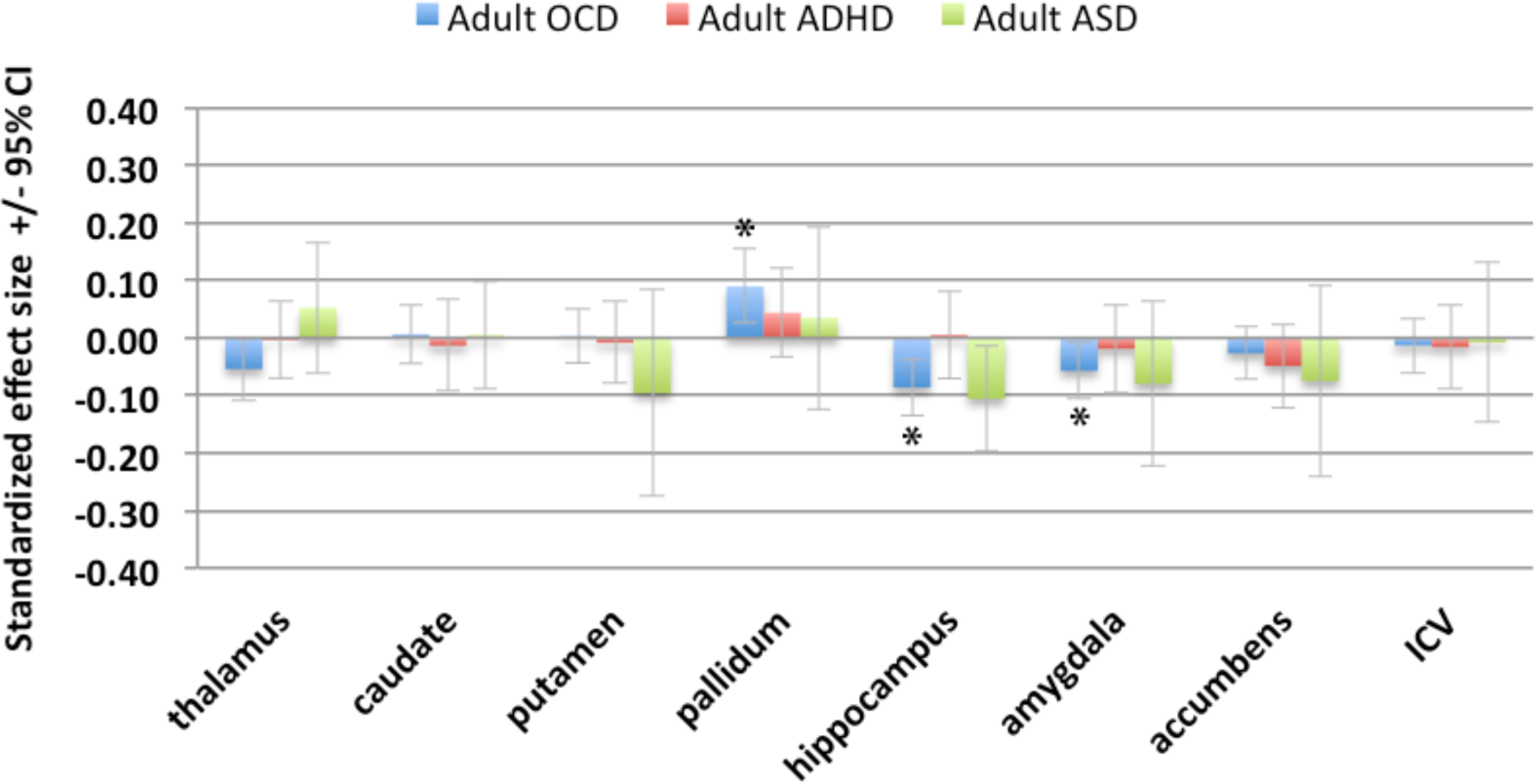
Subcortical volume differences in adults with ADHD, ASD, or OCD compared to controls. Significant results (FDR q ≤ 0.05) are indicated by an asterisk (Supplementary Table S10). For Effect size values across disorders see Supplementary Table S20. Abbreviations: Confidence Interval (CI); Intracranial volume (ICV)

**Figure 3:**
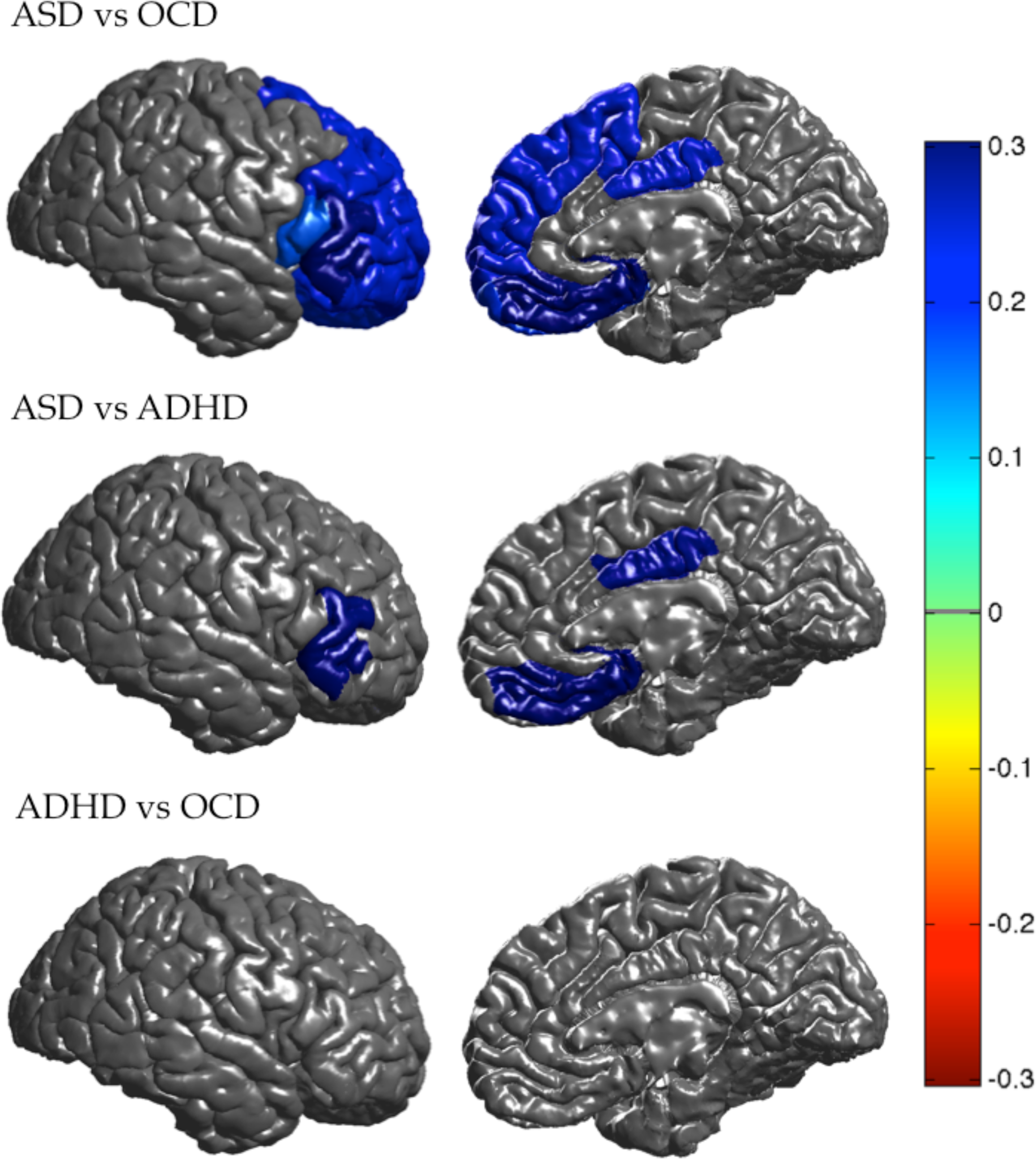
Thicker cortices of several frontal regions in adults with ASD compared to those with OCD or ADHD. Regions that showed a significant (FDR q ≤ 0.05) difference in cortical thickness among adults with ASD, ADHD or OCD. Positive effect sizes *d* (blue) indicate thicker cortices in adults with ASD patients compared to those with ADHD or OCD.

### Influence of medication on cross-disorder effects

Medication status information was incomplete. Table 1a-c lists the numbers of patients for whom information about medication status at the time of scanning was available.

#### Children

The smaller ICV between children with ADHD and those with OCD (effect size=- or those with ASD (effect size=-0.19) may be driven by the unmedicated children (Supplementary Table S23) since ICV did not significantly differed among disorders when comparing the medicated children (Supplementary Tables S24). No *cortical* differences survived multiple comparison correction when comparing unmedicated children among disorders (Supplementary Tables S25 and S26).

Medicated children with OCD had larger amygdala volumes than medicated children with ADHD (effect size=0.43) (Supplementary Table S24). Medicated children with ASD showed a thicker cuneus cortex (effect size=0.60) compared to medicated children with OCD and a thinner middle temporal gyrus (effect size=-0.44) compared to medicated children with ADHD (Supplementary Table S27). No differences in *surface area* differences survived multiple comparison correction when comparing medicated children among disorders (Supplementary Tables S28).

#### Adolescents & Adults

Except for significantly larger *surface area* of the parahippocampal gyrus in unmedicated adults with ASD (effect size=0.33) compared to unmedicated adults with ADHD (Supplementary Table S29), no significant *subcortica*l and *cortical* differences survived multiple comparison correction when comparing unmedicated (Supplementary Tables S30–S34) or medicated (Supplementary Tables S35–S40) adults and adolescents among disorders. Details on disease-specific alterations for unmedicated or medicated patients compared to controls can be found in Supplementary Tables S41-S58.

### Adjusting for Individual Differences in IQ

Information about IQ was incomplete. Table 1a-c shows the number of patients for whom IQ scores were available. We did not have sufficient IQ data to include *adult* patients with OCD into the analysis (Table 1a). Therefore, results for *adults* are based on ASD, ADHD, and HC only.

Adjusting for IQ resulted in similar findings as the main results across all age-groups (Supplementary Tables S59-S67). However, *subcortical volume* analysis did not show smaller hippocampal volumes in children with ADHD and children with ASD compared to those with OCD (Supplementary Table S59). *Cortical thickness* analysis additionally revealed significant thicker cortices of pars orbitalis (effect size=0.20), superior frontal gyrus (effect size=0.22), and frontal pole (effect size=0.23) in adults with ASD compared to adults with ADHD (Supplementary Table S67). Details on disease-specific alterations compared to healthy controls adjusted for IQ can be found in Supplementary Tables S65-S73.

## Discussion

This study comprised the largest neuroimaging investigation to date of structural brain alterations across ADHD, ASD and OCD. Results revealed differing patterns of subcortical and cortical alterations among the disorders across childhood, adolescence, and adulthood. We found ADHD-specific smaller ICV in children and adolescents, and ASD-specific thicker frontal cortices in adults. We did not find OCD-specific alterations across the different age-groups. No brain alterations were shared among all three disorders.

Previous ENIGMA disease working group results, comparing patients with distinct disorders to controls, were mostly replicated, albeit not always using an FDR-corrected threshold. The current study included more patients and considerably more controls than the previously published working group studies (13-17). Accordingly, the present investigation may more accurately represent the normal heterogeneity in the control population. Importantly, our method allowed different mean control group outcomes per cohort, meaning that it statistically accounted for the heterogeneity amongst controls from different cohorts (24).

Overall, results were subtle with small to moderate effect sizes. These effect sizes emerge even after combining dozens of different scanner types and rise above the noise. Large-scale studies like those of the ENIGMA consortium convey another important message mainly by not replicating the extremely large effect sizes that have been found in previous research with smaller samples. Small clinical samples are often rather homogeneous samples carefully selected on the basis of a specific set of in- and exclusion criteria. Homogeneous samples can increase statistical power to discover larger effect sizes, but are typically not representative of the broader population, and such effect sizes are less likely to generalize to the population where patient groups are highly heterogeneous. Although structural brain alterations only indirectly reflect putative alterations at the molecular level, even subtle alterations may be relevant from a pathophysiological perspective (28).

Smaller amygdala volume and thinner frontal and temporal cortices might be shared alterations in children with ASD and ADHD (Supplementary Information SI2). We did not observe similar shared alterations in the adolescents and adults with ASD and ADHD. These findings may be indicative of a more general delayed brain development (18,29). Smaller hippocampus volume might be a shared alteration in adults with ASD and OCD (Supplementary Information SI2). Hippocampal alterations are also described in other psychiatric disorders, such as major depressive disorder, schizophrenia and bipolar disorder (30,31). Decreased hippocampal volume may reflect a disorder non-specific effect, potentially related to chronic stressors (32).

Deficits in social communication and interaction are hypothesized to be related to a thinner temporal cortex (33). Our results fit with the involvement of the temporal cortex in ASD compared to controls, but we did not detect temporal cortex alterations in patients with ASD compared to those with ADHD or OCD. A thicker cortex of several frontal regions was specific to patients with ASD and has been linked to impaired cognitive control and executive dysfunction (13,34). The pattern of thinner temporal and thicker frontal cortices in patients with ASD has been reported in longitudinal studies and suggests accelerated expansion in early childhood, accelerated thinning in later childhood and adolescence, and decelerated thinning in adulthood (35). Although executive dysfunction is present in all three patient groups (4,5), diagnostic categories might differ in executive functioning profiles. Future studies, such as the COMPULS study (36), that focus on neural correlates of executive functioning in all three patient groups will give more insight in this.

Inattention, hyperactivity and impulsivity are the main symptoms of ADHD, presumably modulated by abnormal fronto-striatal circuits (37). Our study confirms frontal surface area and striatal volume alterations in children with ADHD compared to controls, but we did not detect these fronto-striatal alterations in patients with ADHD compared to those with ASD or OCD. Smaller ICV did appear specific to children and adolescents with ADHD. These results support the hypothesis that alterations in ADHD may be due to a delay in brain maturation (29), which possibly normalizes in adulthood. These results are also in line with the genetic correlation between risk for ADHD and smaller ICV (38).

Children with ASD (Supplementary Info SI2) and ADHD seemed to have smaller hippocampal volumes compared to children with OCD. This effect was not detected when adjusting for IQ. Although the sensitivity analysis adjusting for IQ was performed in a smaller subgroup, these findings indicate that the hippocampal volume differences may be driven by IQ differences among patient groups. Indeed, previous studies have shown an association between IQ and hippocampal volume (39). Further cross-disorder analyses adjusted for IQ revealed similar results as the main analyses across all age-groups.

Cross-disorder main effects were not detected when comparing medicated patients and unmedicated patients separately. However, these analyses may have been underpowered to detect the small effect sizes we observed in the larger combined group due to smaller sample sizes when stratifying patients according to medication status.

Two studies performed VBM meta-analyses and reported shared alterations and disease specific alterations between patients with ASD and OCD, and patients with ADHD and OCD, respectively (21,22). Our findings do not corroborate with these findings. This inconsistency might reflect reporting bias in these meta-analyses of published studies and/or differences in analytical methods. FreeSurfer segments brain regions based on probabilistic information from a predefined atlas compared to VBM’s voxel-wise registration. The differences in these methodological approaches may lead to diverging results. Mainly global or regional differences in structure can be inferred from atlas-based FreeSurfer analyses, as opposed to voxel-level morphology with VBM. Thus local morphological alterations may not be detected when averaging across regions (40).

### Strengths and Limitations of the study

This study has several strengths and limitations. As the largest mega-analysis to date, sample size is an obvious strength. Another strength is harmonization of segmentation protocols across all participating sites, reducing variation caused by differences in methods. However, a key limitation is the variation attributable to different scanners and acquisition protocols across cohorts. This issue was mitigated by the formal consideration of potential site differences in all statistical analyses.

Another strength of the study was the use of mega- as opposed to meta-analysis. The comprehensive evaluation of missing data and greater flexibility in control of confounds at the level of individual patients and specific studies are significant advantages. Mega-analyses are also recommended as they avoid the assumptions of within-study normality and known within-study variances, which are especially problematic when including small samples. Indeed our recent study comparing meta- and mega-analytical methods showed that the mega-analytical framework appears to be the better approach for investigating structural neuroimaging data in multi-center studies (24).

We did not perform stratified analyses for reported sex even though ADHD and ASD have a strong sex bias. This issue was mitigated by adjusting for sex in all statistical analyses. Moreover the independent working groups did not observed sex specific effects in their patient groups (13-17).

We chose to differentiate children, adolescents, and adults; cut-offs might not have been optimal, given different disorder onsets. Our rationale was to minimize differences in average age among disorders – in addition to age as a nuisance covariate – and thus to minimize the detection of age effects rather than disease effects. Separate analysis by age group also avoids the difficulties in modeling possibly complex – yet unknown, a priori – nonlinear age effects that might also differ among groups. The primary focus of this manuscript was cross-disorder comparisons. Yet such analyses of age effects are of great interest and should be addressed in future research using multivariate pattern recognition e.g., the support vector machine that can detect informative patterns in the data that may not be identified by traditional linear analyses.

Structural differences among disorders seemed to be minimally affected by medication use and IQ. Nonetheless, we did not have data on medication use and IQ for all patients, indicating insufficient statistical power to address this issue with confidence. We also lacked detailed information on psychotropic treatment. Further efforts are required to draw valid conclusions on the impact of psychotropic medication use on brain structure.

Effects of comorbidity or general phenotypic overlap among ADHD, ASD, and OCD could not be analyzed, because this was not systematically addressed across the cohorts of the different working groups. Presence of comorbidities might have reduced disorder-specific findings. However, excluding comorbid conditions would have ignored complex interactions that are often integral to the disorder. Future studies should test to what extent the comorbid cases differ from the “pure” disorders. Greater consideration of how data may be used in international collaborations such as ENIGMA may influence the collection of data in future studies, which may increase their impact beyond their primary focus.

## Conclusion

To conclude, we found subcortical and cortical differences across different age categories among ADHD, ASD and OCD. We found ASD-specific cortical thickness alterations in the frontal cortex of adult patients and ADHD-specific subcortical alterations in children and adolescents. We did not find shared alterations among the three disorders and shared alterations across any two disorders did not survive multiple comparison corrections. Further work, e.g., multivariate pattern recognition analyses and normative modeling incorporating neural correlates, cognitive and genetic variables will be useful in understanding the mechanisms underlying distinct and shared deficits in these neurodevelopmental disorders.

## Disclosures and acknowledgements

### ADHD working group

#### Disclosures

*David Coghill* served in an advisory or consultancy role for Lilly, Medice, Novartis, Oxford outcomes, Shire and Viforpharma. He received conference support or speaker’s fee by Janssen McNeil, Lilly, Medice, Novartis, Shire and Sunovian. He is/has been involved in clinical trials conducted by Lilly & Shire. The present work is unrelated to the above grants and relationships. *Jonna Kuntsi* has given talks at educational events sponsored by Medice; all funds are received by King’s College London and used for studies of ADHD. *Anders Dale* is a Founder of CorTechs Labs, Inc. He serves on the Scientific Advisory Boards of CorTechs Labs and Human Longevity, Inc., and receives research funding through a Research Agreement with General Electric Healhcare. *Paulo Mattos* was on the speakers’ bureau and/or acted as consultant for Janssen-Cilag, Novartis, and Shire in the previous five years; he also received travel awards to participate in scientific meetings from those companies. The ADHD outpatient program (Grupo de Estudos do Déficit de Atenção/Institute of Psychiatry) chaired by Dr. Mattos has also received research support from Novartis and Shire. The funding sources had no role in the design and conduct of the study; collection, management, analysis, or interpretation of the data; or preparation, review, or approval of the manuscript. *Tobias Banaschewski* served in an advisory or consultancy role for Actelion, Hexal Pharma, Lilly, Lundbeck, Medice, Neurim Pharmaceuticals, Novartis and Shire. He received conference support or speaker’s fee by Lilly, Medice, Novartis and Shire. He is/has been involved in clinical trials conducted by Shire & Viforpharma. He received royalities from Hogrefe, Kohlhammer, CIP Medien, Oxford University Press. The present work is unrelated to the above grants and relationships. *Katya Rubia* received speaker’s fees from Shire, Medice and a grant from Shire pharmaceuticals for another project. *Jan Haavik* has received speaker fees from Eli Lilly, Shire, Medice and Biocodex. *Stephen V. Faraone*, in the past year, received income, potential income, travel expenses continuing education support and/or research support from Tris, Otsuka, Arbor, Ironshore, Shire, Akili Interactive Labs, VAYA, Ironshore, Sunovion, Supernus and Genomind. With his institution, he has US patent US20130217707 A1 for the use of sodium-hydrogen exchange inhibitors in the treatment of ADHD. *Joseph Biederman* is currently receiving research support from the following sources: AACAP, Feinstein Institute for Medical Research, Food & Drug Administration, Genentech, Headspace Inc., Lundbeck AS, Neurocentria Inc., NIDA, Pfizer Pharmaceuticals, Roche TCRC Inc., Shire Pharmaceuticals Inc., Sunovion Pharmaceuticals Inc., and NIH. Dr. Biederman has a financial interest in Avekshan LLC, a company that develops treatments for attention deficit hyperactivity disorder (ADHD). His interests were reviewed and are managed by Massachusetts General Hospital and Partners HealthCare in accordance with their conflict of interest policies. Dr. Biederman’s program has received departmental royalties from a copyrighted rating scale used for ADHD diagnoses, paid by Bracket Global, Ingenix, Prophase, Shire, Sunovion, and Theravance; these royalties were paid to the Department of Psychiatry at MGH. In 2019, Dr. Biederman is a consultant for Akili, Jazz Pharma, and Shire. Through MGH corporate licensing, he has a US Patent (#14/027,676) for a non-stimulant treatment for ADHD, and a patent pending (#61/233,686) on a method to prevent stimulant abuse. In 2018, Dr. Biederman was a consultant for Akili and Shire. *Kerstin Konrad* received speaking fees from Medice, Lilly and Shire. *Josep-Antoni Ramos-Quiroga Josep-Antoni Ramos-Quiroga* was on the speakers’ bureau and/or acted as consultant for Eli-Lilly, Janssen-Cilag, Novartis, Shire, Lundbeck, Almirall, Braingaze, Sincrolab, Medice and Rubió in the last 5 years. He also received travel awards (air tickets + hotel) for taking part in psychiatric meetings from Janssen-Cilag, Medice, Rubió, Shire, and Eli-Lilly. The Department of Psychiatry chaired by him received unrestricted educational and research support from the following companies in the last 5 years: Eli-Lilly, Lundbeck, Janssen-Cilag, Actelion, Shire, Ferrer, Oryzon, Roche, Psious, and Rubió. *Klaus-Peter Lesch* served as a speaker for Eli Lilly and received research support from Medice, and travel support from Shire, all outside the submitted work. *Jan Buitelaar* has been in the past 3 years a consultant to / member of advisory board of / and/or speaker for Janssen Cilag BV, Eli Lilly, Medice, Shire, Roche, and Servier. He is not an employee of any of these companies, and not a stock shareholder of any of these companies. He has no other financial or material support, including expert testimony, patents, royalties. *Barbara Franke* has received educational speaking fees from Shire and Medice. *Susanne Walitza* has received lecture honoraria from Eli-Lilly, Opopharma in the last five years. *Daniel Brandeis* serves as an unpaid scientific consultant for an EU-funded neurofeedback trial. *Georgii Karkashadze* received payment for the authorship of the article and speaker fees from Sanofi and from Pikfarma. *Mario Louza* was on the speakers’ bureau and/or acted as consultant for Janssen-Cilag and Shire in the previous five years; he also received travel awards to participate in scientific meetings from those companies. The present work is unrelated to the above grants and relationships. *Mark Bellgrove* has received speakers fees and travel expenses from Shire within the last 5 years. He is on the Scientific Advisory Board of Novita Healthcare. He is President of the Australian ADHD Professionals Association (AADPA). All other authors from the ENIGMA ADHD working group have no conflicts of interest related to this study.

#### Grant support

*ENIGMA:* received funding from the National Institutes of Health (NIH) Consortium grant U54 EB020403, supported by a cross-NIH alliance that funds Big Data to Knowledge Centers of Excellence (BD2K). We also are supported by the European College for Neuropsychopharmacology (ECNP) by a grant for the ECNP Network ADHD across the lifespan. *ADHD-WUE:* Data collection and analysis was supported by the Deutsche Forschungsgemeinschaft (KFO 125, TRR 58/A1 and A5, SFB-TRR 58/B01, B06 and Z02, RE1632/5-1) and the research leading to these results also received funding from the European Union’s Seventh Framework Programme for research, technological development and demonstration under grant agreement no 602805 (Aggressotype) and the Horizon 2020 research and innovation programme under Grant no. 728018 (Eat2beNICE). *ADHD-DUB1 and DUB2:* The ADHD-DUB1 and DUB2 studies received funding from the Health Research Board Ireland. *ADHD-Mattos:* Ivanei Bramati, Paulo Mattos and Fernanda Tovar-Moll were supported by an IDOR intramural grant. *ADHD200-KKI:* We would like to acknowledge Lindsey Koenig, Michelle Talley, Jessica Foster, Deana Crocetti, Lindsey MacNeil, Andrew Gaddis, Marin Ranta, Anita Barber, Mary Beth Nebel, John Muschelli, Suresh Joel, Brian Caffo, Jim Pekar, Stacy Suskauer. Research was made possible due to the following funding sources: The Autism Speaks Foundation and NIH (R01 NS048527, R01MH078160 and R01MH085328), Johns Hopkins General Clinical Research Center (M01 RR00052), National Center for Resource (P41 RR15241), Intellectual and Developmental Disabilities Research Center (HD-24061) *ADHD200-NYU:* We would like to acknowledge Amy Roy, Andrea McLaughlin, Ariel Schvarcz, Camille Chabernaud, Chiara Fontani, Christine Cox, Daniel Margulies, David Anderson, David Gutman, Devika Jutagir, Douglas Slaughter, Dylan Gee, Emily Brady, Jessica Raithel, Jessica Sunshine, Jonathan Adelstein, Kristin Gotimer, Leila Sadeghi, Lucina Uddin, Maki Koyama, Natan Potler, Nicoletta Adamo, Rebecca Grzadzinski, Rebecca Lange, Samantha Adelsberg, Samuele Cortese, Saroja Bangaru, Xinian Zuo, Zarrar Shehzad and Zoe Hyde. Data collection was made possible thanks to funding from NIMH (R01MH083246), Autism Speaks, The Stavros Niarchos Foundation, The Leon Levy Foundation, and an endowment provided by Phyllis Green and Randolph Cowen. *ADHD200-Peking*: we would like to acknowledge Jue-jing Ren, De-yi Wang, Su-fang Li, Zu-lai Peng, Peng Wang, Yun-yun Zhu, Zhao Qing. Research was made possible due to the following funding sources: The Commonwealth Sciences Foundation, Ministry of Health, China (200802073), The National Foundation, Ministry of Science and Technology, China (2007BAI17B03), The National Natural Sciences Foundation, China (30970802), The Funds for International Cooperation of the National Natural Science Foundation of China (81020108022), The National Natural Science Foundation of China (8100059), Open Research Fund of the State Key Laboratory of Cognitive Neuroscience and Learning. *ADHD200-OHSU:* We would like to acknowledge the Advanced Imaging Research Center, Bill Rooney, Kathryn L. Mills, Taciana G. Costa Dias, Michelle C. Fenesy, Bria L. Thurlow, Corrine A. Stevens, Samuel D. Carpenter, Michael S. Blythe, Colleen F. Schmitt. Research was made possible due to the following funding resources: K99/R00 MH091238 (Fair), R01 MH086654 (Nigg), Oregon Clinical and Translational Research Institute (Fair), Medical Research Foundation (Fair), UNCF/Merck (Fair), Ford Foundation (Fair). *ADHD-UKA:* KFO-112 and IRTG1328 was supported by the German Research Foundation (DFG). *DAT-London:* This work was supported in part by UK Medical Research, Council Grant G03001896 to J Kuntsi and NIH grants, R01MH62873 and R01MH081803 to SV Faraone. *IMpACT:* The IMpACT study was supported by a grant from the Brain & Cognition Excellence Program and a personal Vici grant (to Barbara Franke) of the Netherlands Organization for Scientific Research (NWO, grant numbers 433-09-229 and 016-130-669) and in part by the Netherlands Brain Foundation (grant number, 15F07[2]27)and the BBMRI-NL (grant CP2010-33). Funding was also provided by a pilot grant of the Dutch National Research Agenda for the NeuroLabNL project. The research leading to these results also received funding from the European Community’s Seventh Framework Programme (FP7/2007–2013) under grant agreement no. 602805 (Aggressotype), no. 278948 (TACTICS), and no. 602450 (IMAGEMEND). In addition, the project received funding from the European Union’s Horizon 2020 research and innovation programme under the Marie Sklodowska-Curie grant agreement no. 643051 (MiND), under grant agreement no. 667302 (CoCA) and the grant agreement no. 728018 (Eat2beNICE). *NYU ADHD:* NYU data collection and sharing was supported by NIH grants T32MH67763, R01MH083246, K23MH087770, R01MH094639, and U01MH099059 and a grant from the Stavros S. Niarchos Foundation. *UAB-ADHD:* The study and its contributors received funding from the Ministerio de Economía y Competitividad under research grant SAF2012-32362 and : PI12/01139 from the Department of Health of the Government of Catalonia. Additional funding was obtained from the Generalitat de Catalunya. *ZI-CAPS*: The Neurofeedback study was partly funded by the project D8 of the Deutsche Forschungsgesellschaft collaborative research center 636. *ADHD-Rubia:* The study was funded by the UK Department of Health via the National Institute of Health Research Centre (BRC) for Mental Health South London and the Maudsley NHS Foundation Trust and the Institute of Psychiatry, King’s College London. *CAPS_UZH:* The data contributed to this study were collected in two projects on ADHD and OCD in children and adolescents, supported by the Swiss National Science Foundation (projects No. 136249 Sinergia and No. 320030_130237) and the Hartmann Müller Foundation (No. 1460). *NeuroIMAGE:* This work was supported by NIH Grant R01MH62873, NWO Large Investment Grant 1750102007010 and grants from Radboud University Medical Center, University Medical Center Groningen and Accare, and VU University Amsterdam. This work was also supported by grants from NWO Brain & Cognition (433-09-242 and 056-13-015) and from ZonMW (60-60600-97-193). Further support was received from the European Union FP7 programmes TACTICS (278948) and IMAGEMEND (602450). *MTA:* Data collection and sharing for this project was funded by the NIDA MTA Neuroimaging Study (National Institute on Drug Abuse Grant Contract #: HHSN271200800009C). *NIH:* studies were supported by intramural grants at the National Institute of Mental Health and National Human Genome Research Institute. *OHSU:* The OHSU work was supported by NIMH grants R01MH86654, MH099064, and MH115357. *UCHZ:* This work was supported by the University Research Priority Program “Integrative Human Physiology” at the University of Zurich. *ACPU:* This research was conducted within the Academic Child Psychiatry Unit, University of Melbourne, Royal Children’s Hospital and the Developmental Imaging research group, Murdoch Children’s Research Institute, Melbourne, Victoria. National Health and Medical Research Council of Australia (NHMRC) project grants 384419 and 569533 provided funds for the data collection. It was also supported by the Murdoch Children’s Research Institute, the Royal Children’s Hospital and the Children’s MRI Centre, The Royal Children’s Hospital Foundation, and the RCH Mental Health Service, Department of Paediatrics The University of Melbourne and the Victorian Government’s Operational Infrastructure Support Program. Tim Silk was supported by a NHMRC Career Development Award. *NICAP:* The Neuroimaging of the Children’s Attention Project was funded by the National Medical Health and Research Council of Australia (NHMRC; project grant #1065895). Earlier funding of the Children’s Attention Project was as funded by an NHMRC project grant #1008522 and a grant from the Collier Foundation. This research was conducted within the Developmental Imaging research group, Murdoch Children’s Research Institute and the Children’s MRI Centre, The Royal Children’s Hospital, Melbourne, Victoria. It was supported by the Murdoch Children’s Research Institute, The Royal Children’s Hospital, The Royal Children’s Hospital Foundation, Department of Paediatrics at The University of Melbourne and the Victorian Government’s Operational Infrastructure Support Program. *Tübingen:* The recruitment of the Tübingen sample was funded by the Deutsche Forschungsgemeinschaft (DFG grant: ET 112/5-1). *Dundee:* This work was supported by a grant from TENOVUS SCOTLAND and was conducted in collaboration with the Dundee site of the ADHD Drugs Use Chronic Effects (ADDUCE) study (EU FP7 agreement No. **260576).** *ePOD:* The neuroimaging studies of the ePOD-MPH trial (NTR3103) were supported by faculty resources of the Academic Medical Center, University of Amsterdam, and by grant 11.32050.26 from the European Research Area Network Priority Medicines for Children (Sixth Framework Programme) to Liesbeth Reneman. *Sao Paulo:* The present investigation was supported by a 2010 NARSAD Independent Investigator Award (NARSAD: The Brain and Behavior Research Fund) awarded to Geraldo F. Busatto. Geraldo F. Busatto is also partially funded by CNPq-Brazil. *Sussex:* This study was supported by funding from Brighton and Sussex Medical School and the Dr. Mortimer and Dame Theresa Sackler Foundation. *Clinic Barcelona:* This work has received financial support from two grants, *Fundació la Marató de TV3-2009* (project number: 091810) and *Fondo de Investigaciones Sanitarias*, of the Spanish Ministry of Health (project number: PI11/01419). *Maarten Mennes:* supported by a Marie Curie International Incoming Fellowship within the 7th European Community Framework Programme, grant agreement n° 327340. *Jan Haavik*: Stiftelsen Kristian Gerhard Jebsen (SKGJ-MED 02). *Steve Faraone:* K.G. Jebsen Centre for Research on Neuropsychiatric Disorders, University of Bergen, Bergen, Norway

### ASD working group

#### Disclosures

*Dr. Anagnostou* has served as a consultant or advisory board member for Roche and Takeda; she has received funding from the Alva Foundation, Autism Speaks, Brain Canada, the Canadian Institutes of Health Research, the Department of Defense, the National Centers of Excellence, NIH, the Ontario Brain Institute, the Physicians’ Services Incorporated (PSI) Foundation, Sanofi-Aventis, and SynapDx, as well as in-kind research support from AMO Pharma; she receives royalties from American Psychiatric Press and Springer and an editorial honorarium from Wiley. *Dr. Arango* has served as a consultant for or received honoraria or grants from Acadia, Abbott, Amgen, CIBERSAM, Fundación Alicia Koplowitz, Instituto de Salud Carlos III, Janssen-Cilag, Lundbeck, Merck, Instituto de Salud Carlos III (co-financed by the European Regional Development Fund “A way of making Europe,” CIBERSAM, the Madrid Regional Government [S2010/BMD-2422 AGES], the European Union Structural Funds, and the European Union Seventh Framework Programme under grant agreements FP7-HEALTH-2009-2.2.1-2-241909, FP7-HEALTH-2009-2.2.1-3-242114, FP7-HEALTH-2013-2.2.1-2-603196, and FP7-HEALTH-2013-2.2.1-2-602478), Otsuka, Pfizer, Roche, Servier, Shire, Takeda, and Schering-Plough. *Dr. Freitag* has served as a consultant for Desitin regarding issues on ASD. *Dr. De Martino* is a coauthor of the Italian version of the Social Responsiveness Scale, for which she may receive royalties. *Dr. Rubia* has received speaking honoraria from Eli Lilly, Medice, and Shire. *Dr. Buitelaar* has served as a consultant, advisory board member, or speaker for Eli Lilly, Janssen-Cilag, Lundbeck, Medice, Novartis, Servier, Shire, and Roche, and he has received research support from Roche and Vifor. *Dr. Gallagher* received funding from are the Meath Foundation and The National Childrens Research Centre in Ireland. The other authors report no financial relationships with commercial interests.

#### Grant support

ENIGMA received funding from NIH Consortium grant U54 EB020403 to *Dr. Paul Thompson*, supported by a cross-NIH alliance that funds Big Data to Knowledge Centers of Excellence (BD2K). This research was further supported by the European Community’s Seventh Framework Programme (FP7/2007–2013) under grant agreement number 278948 (TACTICS), and the Innovative Medicines Initiative Joint Undertaking under grant agreement number 115300 (EU-AIMS), resources of which are composed of financial contributions from the European Union’s Seventh Framework Programme (FP7/2007–2013) and the European Federation of Pharmaceutical Industries and Associations companies’ in-kind contribution. The Canadian samples were collected as part of the Province of Ontario Neurodevelopmental Disorders (POND) Network, funded by the Ontario Brain Institute (grant IDS-I l-02 to *Dr. Anagnostou* and *Dr. Lerch*). *Dr Calvo* has received The Marató TV3 Foundation Grant No.091710, the Carlos III Health Institute (PI091588) co-funded by FEDER funds/European Regional Development Fund (ERDF), “a way to build Europe”. *Declan Murphy* has received funding from the Innovative Medicines Initiative 1 and 2 Joint Undertaking under grant agreement no.115300 (EU AIMS) and no. 777394 (AIMS-2-TRIALS), the National Institute for Health Research (NIHR) Biomedical Research Centre at South London and Maudsley NHS Foundation Trust and King’s College London, and a Medical Research Council grant no. G0400061.

### OCD working group

#### Disclosures

*Dr. Baker* has received research support from the National Institute of Mental Health (NIMH) and Valera Health. *Dr. Brennan* has received consulting fees from Rugen Therapeutics and Nobilis Therapeutics and research grant support from Eli Lilly, Transcept Pharmaceuticals, and Biohaven Pharmaceuticals. *Dr. Mataix-Cols* receives royalties for contributing articles to UpToDate, Wolters Kluwer Health and fees from Elsevier for editorial tasks (all unrelated to the submitted work). *Dr Simpson* received research support for multi-site clinical trial from Biohaven, Inc; royalties from UpToDate, Inc and Cambridge University Press. *Dr. Stein* has received research grants and/or consultancy honoraria from AMBRF, Biocodex, Cipla, Lundbeck, National Responsible Gambling Foundation, Novartis, Servier, and Sun in the past 3 years. *Dr. Walitza* has received lecture honoraria Opopharma in the last 3 years. Her work was supported in the last 3 years by the Swiss National Science Foundation (SNF), diverse EU FP7s, HSM Hochspezialisierte Medizin of the Kanton Zurich, Switzerland, Bfarm Germany, Zinep, Hartmann Müller Stiftung, Olga Mayenfisch. All other authors from the ENIGMA OCD working group have no conflicts of interest related to this study.

#### Grant support

The **ENIGMA-Obsessive Compulsive Disorder Working-Group** gratefully acknowledges support from the NIH BD2K award U54 EB020403 (PI: *Dr. Thompson*) and Neuroscience Amsterdam, IPB-grant to *Dr. Schmaal* & *Dr. van den Heuvel*. Supported by the Japan Society for the Promotion of Science (JSPS; KAKENHI Grants No. 18K15523 to *Dr. Abe*, No. 16K19778, No. 18K07608 to *Dr. Nakamae,* No. 16K04344 to *Dr. Hirano*, and No. 26461762 to *Dr. Nakagawa*); Ontario Mental Health Foundation Research Training Fellowship to *Dr. Ameis*; the Fundação de Amparo à Pesquisa do Estado de São Paulo (FAPESP, São Paulo Research Foundation; Grant No. 2011/21357-9; the EU FP7 project TACTICS (Grant No. 278948 to *Dr. Buitelaar*); the National Natural Science Foundation of China (No. 81560233 to *Dr. Cheng*); the International Obsessive-Compulsive Disorder Foundation (IOCDF) Research Award to *Dr. Gruner*; the Dutch Organization for Scientific Research (NWO) (grants 912-02-050, 907-00-012, 940-37-018, and 916.86.038); the Netherlands Society for Scientific Research (NWO-ZonMw VENI grant 916.86.036 to *Dr. van den Heuvel*; NWO-ZonMw AGIKO stipend 920-03-542 to *Dr. de Vries*), a NARSAD Young Investigator Award to *Dr. van den Heuvel*, and the Netherlands Brain Foundation (2010(1)-50 to *Dr. van den Heuvel*); the Federal Ministry of Education and Research of Germany (No. BMBF-01GW0724 to *Dr. Kathmann*); the Alberta Innovates Translational Health Chair in Child and Youth Mental Health and the Ontario Brain Institute funding *Dr. Arnold*; the Deutsche Forschungsgemeinschaft (DFG; Grant No. KO 3744/7-1 to *Dr. Koch*); the Helse Vest Health Authority (No. 911754, 911880 to *Dr. Kvale*) and the Norwegian Research Council (No. HELSEFORSK 243675 to *Dr. Kvale*); the Wellcome Trust and a pump priming grant from the South London and Maudsley Trust, London, UK (Project Grant No. 064846 to *Dr. Mataix-Cols*); the Generalitat de Catalunya (AGAUR 2017 SGR 1247 to *Dr. Menchón*); the PhD-iHES program (FCT fellowship Grant No. PDE/BDE/113601/2015 to Dr. Moreira); the Japanese Ministry of Education, Culture, Sports, Science and Technology (Grant-in-Aid for Scientific Research (C) 22591262, 25461732, 16K10253 to *Dr. Nakao*); the Italian Ministry of Health (No. RC13-14-15-16-17-18A to *Dr. Spalletta, Dr. Piras Fabrizio, Dr. Piras Federica*); the National Center for Advancing Translational Sciences (Grant No. UL1TR000067/KL2TR00069) and the NIMH (Grant No. R33MH107589 & R01MH111794) grants to *Dr. Stern*; the SA MRC funding and the National Research Foundation of South Africa *Dr. Stein and Dr. Lochner*; the Canadian Institutes of Health Research, and British Columbia Provincial Health Services Authority funding *Dr. Stewart*; the Netherlands Organization for Scientific Research (NWO/ZonMW Vidi 917.15.318 to *Dr. van Wingen*); the Wellcome-DBT India Alliance (Grant No. 500236/Z/11/Z to *Dr. Venkatasubramanian*); the National Natural Science Foundation of China (No. 81371340) and the Shanghai Key Laboratory of Psychotic Disorders (No. 13dz2260500) to *Dr. Wang*; the Government of India grants from the Department of Science and Technology (Grants No. SR/S0/HS/0016/2011 to *Prof. Y.C. Janardhan Reddy*, and DST INSPIRE faculty grant -IFA12-LSBM-26 to *Dr. Janardhanan C. Narayanaswamy*) and from the Department of Biotechnology (Grants No. BT/PR13334/Med/30/259/2009 to *Prof. Y.C. Janardhan Reddy*, and No. BT/06/IYBA/2012 to *Dr. Janardhanan C. Narayanaswamy*); the Dana Foundation and NARSAD to *Dr. Fitzgerald*; the NIMH (Grant No. R01MH107419 and K23MH082176 to *Dr. Fitzgerald*; Grant No. R21MH101441 to *Dr. Marsh*; Grant No. R21MH093889 to *Drs. Marsh & Simpson*; Grant No. K23-MH104515 to *Dr. Baker*; Grant No. R01MH081864 to *Drs. O’Neill, Piacentini;* and Grant No. R01MH085900 to *Drs. O’Neill* and *Feusner*); the NIMH and the David Judah Fund at the Massachusetts General Hospital (Grant No. K23-MH092397 to *Dr. Brennan*); the Marató TV3 Foundation (Grants No. 01/2010 and 091710 to *Dr. Lazaro*); the Carlos III Health Institute (PI040829 to *Dr Lazaro*, Grant CPII16/00048 and Project Grants PI13/01958 & PI16/00889 to *Dr. Soriano-Mas*), co-funded by FEDER funds/European Regional Development Fund (ERDF), a way to build Europe; the AGAUR (2017 SGR 881 to *Dr. Lazaro*); the Michael Smith Foundation for Health Research funding *Dr. Jaspers-Fayer* and *Dr. Stewart*; the Carlos III Health Institute (PI14/00419 to *Dr. Alonso*, Grant No. FI17/00294 to Ignacio Martínez-Zalacaín, Project Grant No. PI16/00950 to *Dr. Menchón*); and the Japan Agency for Medical Research and Development (AMED Grant Number JP18dm0307002 to *Dr. Hirano*).

